# Evaluating Wayfinding Designs in Healthcare Settings through EEG Data and Virtual Response Testing

**DOI:** 10.1101/2021.02.10.430638

**Authors:** Saleh Kalantari, Vidushi Tripathi, James D. Rounds, Armin Mostafavi, Robin Snell, Jesus G. Cruz-Garza

**Affiliations:** Department of Design and Environmental Analysis, Cornell University, Ithaca, NY, United States; Human Development, Cornell University, Ithaca, NY, United States; Institute of Architectural Sciences, Vienna University of Technology, Vienna, Austria; Parkin Architects, Toronto, Canada

**Keywords:** Wayfinding, Healthcare Design, Virtual Reality, EEG, Spatial Behavior

## Abstract

Wayfinding difficulties in healthcare facilities have been shown to increase anxiety among patients and visitors and reduce staff operational efficiency. Wayfinding-oriented interior design features have proven beneficial, but the evaluation of their performance is hindered by the unique nature healthcare facilities and the expense of testing different navigational aids. This study implemented a virtual-reality testing platform to evaluate the effects of different signage and interior hospital design conditions during navigational tasks; evaluated through behavioral responses and mobile EEG. The results indicated that using color to highlight destinations and increase the contrast of wayfinding information yielded significant benefits when combined with wayfinding-oriented environmental affordances. Neural dynamics from the occipital cortex showed beta-band desynchronization with enhanced color condition and additional theta-band desynchronization with enhanced environmental affordance. This multimodal testing platform has the potential to establish a robust body of evidence for future wayfinding design strategies.

## INTRODUCTION

Wayfinding systems—signs, color-schemes, and similar features intended to aid in navigation—can strongly contribute to the comfort and wellbeing of patients, visitors, and staff members in healthcare settings (Carpman et al., 1990; Foxall & Hackett, 1994; Nelson-Shulman, 1983; Ulrich et al., 2008). Unfortunately, these design features are frequently treated as an “afterthought” rather than as an integral part of the architectural design process in healthcare facilities (Devlin, 2014). In a study conducted in 1990, the annual cost of maintaining the ad-hoc wayfinding system in a tertiary-care hospital was calculated to be more than $220,000 per year ($448 per bed per year), and the researchers found that direction-giving by hospital employees other than information staff occupied more than 4,500 staff hours per year (Zimring, 1990). Difficulties in wayfinding due to inadequate design features have been shown to be a significant source of stress for hospital patients (Devlin, 2014), as well as a significant burden on hospital employees and an obstacle to operational efficiency (Peponis et al., 1990).

There is a growing body of research in wayfinding that investigates the most effective cues for helping people find a path through the physical environment (Apelt et al., 2007; Calori & Vanden-Eynden, 2015; Gibson, 2009; Levine, 1982; Levine et al., 1984; Ophir et al., 2009; Pollet & Haskel, 1979; Rodrigues et al., 2019; Sharma et al., 2017; Weisman, 1981). However, empirical research in wayfinding that focuses specifically on healthcare settings is limited; and in general, logistical factors make it very difficult to carry out any kind of rigorous, comparative studies of wayfinding features in the context of constructed facilities (Ulrich et al., 2008). Some of the most frequently used wayfinding design strategies in healthcare settings include developing a distinct color scheme for each unit, adding prominent pictograms and recognizable icons, and adjusting architectural features to highlight destinations and facilitate move/stop behavior (Carpman, 1993; Huelat, 2004; Kalantari & Snell, 2017; Passini et al., 2000; Rooke et al., 2010; Ulrich et al., 2008). The relative effectiveness of these different design interventions has not been empirically tested to any great extent. Furthermore, the utility of any given wayfinding strategy is likely dependent on the manner in which it is implemented in the overall design of a particular building, which means that the small number of existing research studies in this area may not be readily generalizable to new facility designs.

The current project addressed these issues by developing an effective method that can be used during the design development process to test and optimize wayfinding design strategies in a specific acute care hospital facility. The goal was to evaluate the ease of wayfinding in specific facility designs, catch potential problems prior to construction, and enable evidence-gathering to support (or refute) more innovative and creative wayfinding design strategies. This was accomplished through the use of an immersive high-resolution virtual reality (VR) platform, which allowed the study of participant behavioral and neural responses during wayfinding tasks under different design conditions. Using the VR approach makes it possible to simply switch out different signs, colors-patterns, and other wayfinding features (without incurring any construction costs), and thereby evaluate the likely success of these designs for hospital users and tweak them to remove problem spots.

There are some limitations in the use of VR for behavioral studies, since the bodily motions, sensory immersion, and physiological responses to VR contexts may not precisely mirror real-world experiences. However, the alternative to using VR testing of wayfinding strategies in healthcare design is to conduct *no testing at all*, which is currently the standard practice in hospital wayfinding systems. Due to the unique nature of each healthcare facility, as well as the tremendous expense and logistical implausibility of evaluating different wayfinding solutions in an occupied real-world building, there is currently no good method of rigorously analyzing the relative success of different wayfinding design strategies. Our proposed platform will allow designers to advance beyond pure speculation or anecdotal evidence, and to instead collect detailed and objective feedback about proposed wayfinding features. This approach follows a general call in the design field for the greater use of VR studies as a form of pre-construction testing (Hölscher et al., 2006; Jansen-Osmann, Schmid, & Heil, 2007; Jansen-Osmann & Fuchs, 2006; Jansen-Osmann & Wiedenbauer, 2004; Tang et al., 2009; Kalantari, 2016; Kalantari & Neo, 2020; Werner & Schindler, 2004). Another advantage of the VR setting is that it allows for the use of physiological sensors that would be extremely unwieldy or distracting (or simply technologically infeasible) to use in a real-world testing environment. The use of physiological data creates an additional layer of empirical feedback about human responses to designs (Cruz-Garza et al. 2021). Part of the motivation for the current project was to introduce more objective physiological data in evaluating participant responses during wayfinding tasks, by applying electroencephalography (EEG), electrocardiography (ECG), galvanic skin response (GSR), and motion-tracking technology. The collected physiological data can be triangulated against the participants’ subjective feedback, and synchronized chronologically to their actions and movements within the VR environment. For example, we can evaluate whether or not a participant looked at a particular sign, how long they looked at it, any changes in their heart rate or electrical brain activity while looking at the sign, how quickly and accurately they then moved toward the target destination, as well as their conscious feedback about the sign’s effectiveness. In the current research we conducted a study to evaluate different color patterns, signage enhancements (e.g., pictograms), and architectural features and queues for wayfinding in a specific healthcare facility design-build project. In this work, we also emphasized the creation of a streamlined data collection protocol based on the VR and psychophysiological data that can easily collect and synchronize participant data and thereby empower other researchers and designers to use this approach in evaluating their own wayfinding design strategies in various buildings.

## OVERVIEW OF PRIOR WAYFINDING RESEARCH IN HEALTHCARE SETTINGS

Wayfinding in healthcare facilities can be a challenging task. Hospitals are large and complex, and most people are unfamiliar environments (Devlin, 2014; Mollerup, 2009). Furthermore, spatial issues often develop and/or are exacerbated over time as hospitals are renovated and new additions are built (Cheng & Pérez-Kriz, 2014; Mollerup, 2009; Rousek & Hallbeck, 2011). The population that has to navigate through these complicated building typically includes a large number of first-time and infrequent visitors, as well as individuals who may be in a temporary state that impairs their judgment, perception, or mobility (e.g., from sickness, anxiety, injury, etc.).

Prior studies on wayfinding in hospital environments have sought to identify disparate features that may help or hinder such individuals in finding their destinations. Unfortunately, this literature is limited and there are very few robust post-occupancy studies evaluating hospital wayfinding designs. The expense of conducting such studies, along with the unique nature of each facility, makes it unlikely that extensive rigorous comparative studies of hospital wayfinding strategies will ever be carried out in real-world contexts (Kalantari & Snell, 2017). Based on the limited and often anecdotal evidence that does exist, researchers and designers have gravitated toward a handful of features that are believed to influence wayfinding success in hospital settings, including landmarks (Passini, 1984; Scialfa et al., 2004; Weisman, 1981), architectural features such as color, texture, and room height for defining different areas and creating sight-lines (Ahn, 2006; Carpman et al., 1985; Hölscher et al., 2013), visual arrangements of signs and furniture (O’Neill, 1991), logical floor-plans (Baskaya et al., 2004; Passini, 1984; Slone et al., 2015; Weisman, 1981), or some combination of the above (Carpman, 1993; Huelat, 2004; Passini et al., 2000; Pati et al., 2015; Rooke et al., 2010; Ulrich et al., 2008).

The overall results from this literature provisionally indicate that wayfinding success in hospitals may be linked both to the structural features of buildings (floor-plans, room shapes, sightlines) and to overlaid directional cues (landmarks, signs, color schemes) (Devlin, 2014; Marquardt, 2011). The mutual impact of these factors is reminiscent of Carpman and colleagues’ (1985) influential view that architectural design is a combination of environmental affordances (what the structure suggests can occur) and manifest cues (overlaid indications of what should occur). An interesting hospital design study by Vilar and colleagues (2014) placed environmental affordances in terms of corridor width and brightness in direct competition with manifest cues—signs that directed participants in a non-intuitive direction. The results indicated that visitors in emergency situations tended to rely more strongly on the manifest cues, whereas those arriving for more mundane tasks were more likely to attend to the environmental affordances. Ideally, of course, healthcare designs should seamlessly merge structural features and overt cues to enable successful wayfinding, but this research direction should remind us to consider the needs and responses of users who may have different levels of urgency in their tasks as well as different visual and physical capabilities. While healthy, unharried participants in a study context may respond well to subtle environmental cues, researchers should remember that other hospital staff and visitors may have different needs and experiences.

## MATERIALS AND METHODS

### Facility Design Conditions and Study Hypotheses

The researchers’ immediate goal was to carry out a study on the success of wayfinding design strategies, using a healthcare facility during design by our industry partner as the testing environment. The healthcare facility was the Corner Brook Acute Care Hospital, located in Corner Brook, Canada was in the design development phase during this study. Two specific parts of this large hospital complex were selected to be used in the study, and design details about those portions of the facility were imported into our virtual testing platform. Drawing from the prior literature of healthcare wayfinding design— particularly the concepts of environmental affordances and manifest cues (Devlin, 2014)—we adjusted the virtual environment to create three distinct design conditions:

- Condition A. Baseline: standard signage and minimal environmental contrast.
- Condition B. Added color to highlight destinations and to increase the contrast of wayfinding information against the overall environment.
- Condition C. Includes the color from Condition B, along with added architectural features and textures to further highlight destinations, prompt or deter movement, signal functional changes between different areas of the building, and added distinguishable patterns to wayfinding signs.

It is important to note that this study was not designed to obtain definitive conclusions about the value of color, graphics, and architectural feature, particularly because we examined only one possible implementation of these design principles. As discussed in the previous section, the effectiveness of a wayfinding strategy very likely depends on how it is implemented in a specific building, and it may also be affected by a variety of confounding/external variables (user demographics, user capabilities, task urgency, etc.). Our goal was rather to collect specific evidence about the effectiveness of proposed wayfinding designs in this particular facility in a convenience sample (primarily undergraduate students, from an entirely different geographic area than that of the planned hospital location), and to demonstrate the exciting potential of VR immersion with physiological sensors as a method of pre-construction testing (**Figures 1 and 2**). In the future, with more widespread adoption of this testing approach for many individual projects and population groups, a more comprehensive corpus of empirical findings on wayfinding design strategies may begin to emerge.

**Figure 1.**
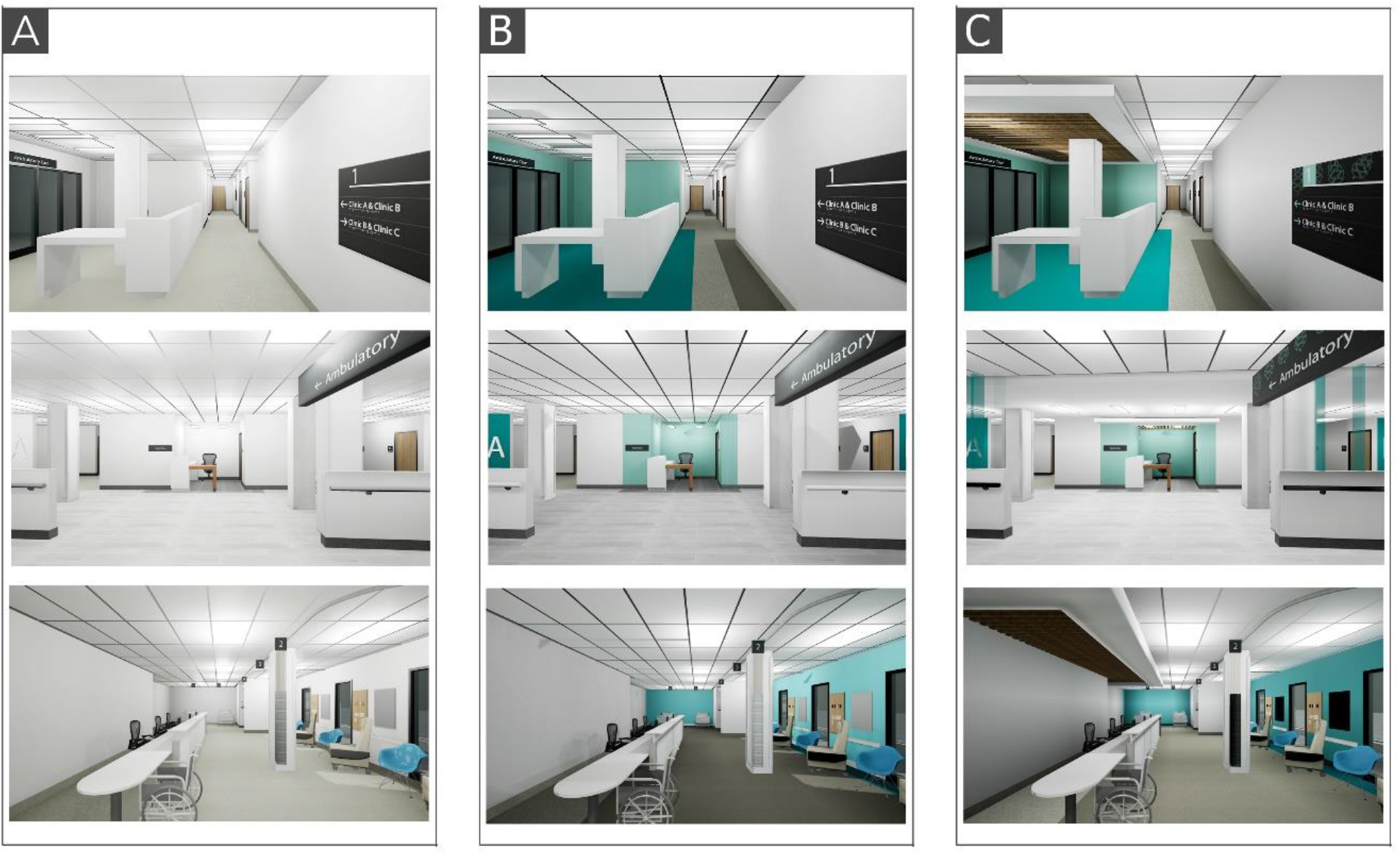
Screen-captures from the VR environment showing comparative examples of the wayfinding design features in the Baseline Condition (A), the Enhanced Color Condition (B), and the Enhanced Color, Graphics, and Architectural Feature Condition (C).

**Figure 2.**
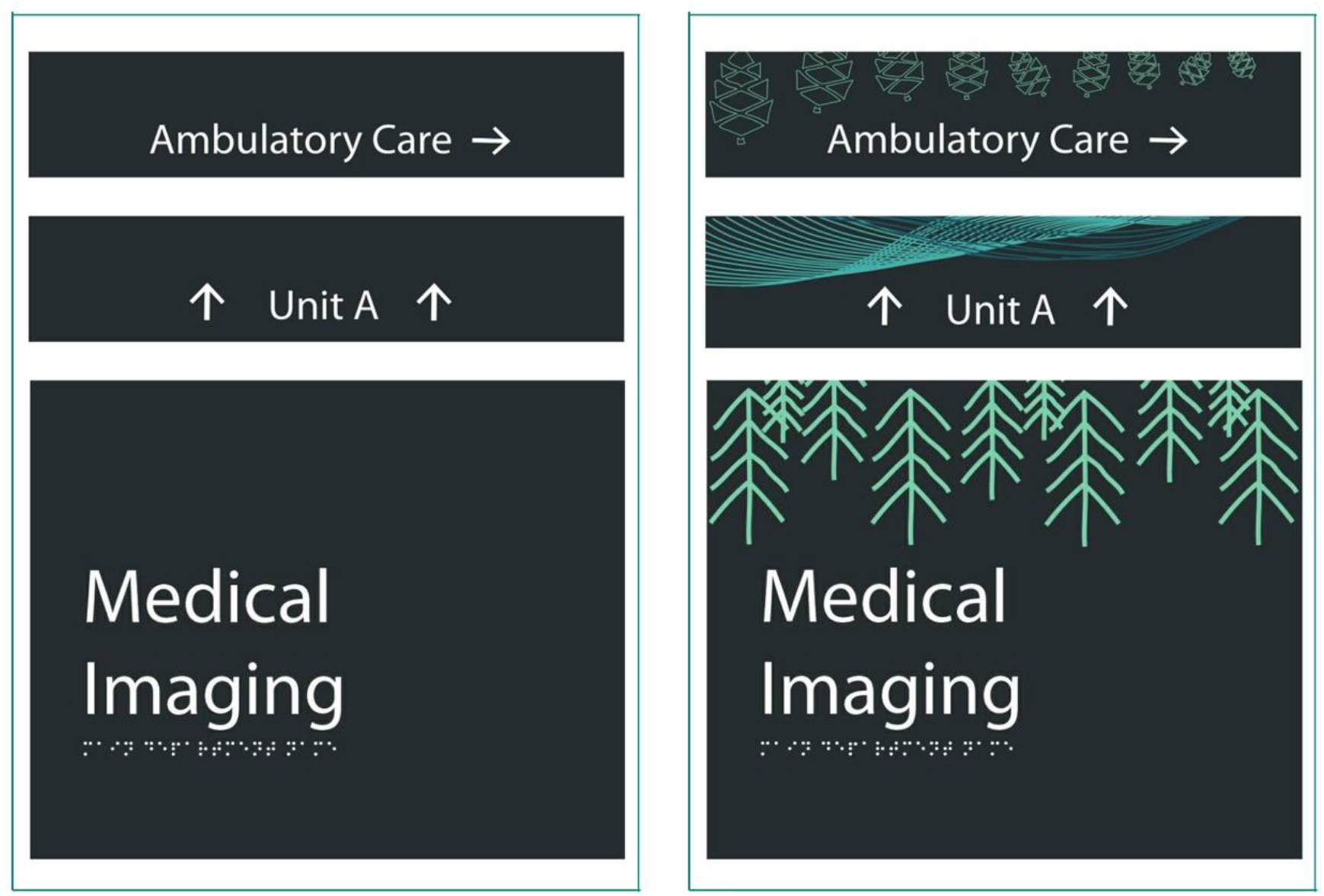
Example of the thematic graphics and contrast that were added in design conditions C (at right), compared to the signage in design condition A and B (at left).

Participants in the study were randomly assigned to one of the three design conditions and were asked to complete a series of wayfinding tasks in the VR environment while behavioral and physiological data was collected synchronously. The data were statistically compared between the three design conditions to test the study hypotheses:

**H1:** Behavioral metrics related to successful wayfinding will be significantly higher among participants who complete the tasks in the hospital design with enhanced color (Condition B), compared to the baseline design (Condition A). These behavioral metrics include:

H1a: Lower self-reported mental fatigue, stress, and confusion.
H1b: Less time required for wayfinding task completion.
H1c: More efficient and accurate orientation behavior and the correct choice of direction.
**H2:** Behavioral metrics related to successful wayfinding will be significantly higher among participants who complete the tasks in the hospital design with enhanced color, graphics, and architectural feature condition (Condition C), compared to the other two design conditions (Conditions A and B). These behavioral metrics include:

H2a: Lower self-reported mental fatigue, stress, and confusion.
H2b: Less time required for wayfinding task completion.
H2c: More efficient and accurate orientation behavior and the correct choice of direction.
**H3:** Neural patterns associated with spatial awareness and navigational processing, e.g., alpha, beta, and theta band EEG in parietal and occipital areas will be heightened when participants complete the tasks in the hospital design with enhanced color condition (Condition B), compared to the baseline design (Condition A).
**H4:** Neural patterns associated with spatial awareness and navigational processing, e.g., alpha, beta, and theta band EEG in parietal and occipital areas, will be heightened when participants complete the tasks in the hospital design with enhanced color, graphics, and architectural feature (Condition C), compared to the other two design conditions (Conditions A and B).

### Virtual Reality Development

The creation of the virtual hospital environment was carried out by importing architectural design documents into Epic Games’ Unreal Engine (www.epicgames.com). Most of the modeling and UV-mapping took place within Autodesk 3ds Max (www.autodesk.com). The Unreal Engine uses Blueprint scripting, allowing for a quick learning curve on the part of researchers and designers who may want to expand or replicate our work. All of the front-end interaction and user interactivity in our testing environment also leverages the Blueprint platform.

The camera height was fixed at 1.70 m (corresponding to the average human eye-height), but participants were otherwise allowed to move freely throughout the environment and to alter the camera angle (looking up or down). Custom interactive widgets allowed self-reported Likert-scale responses to be collected directly in the VR environment. The hospital designs were presented to study participants using an HTC Vive Pro head-mounted display, through a gaming desktop with a resolution of 1440×1600 pixels. The Vive Pro provides a 110-degree horizontal field of view with a 90 Hz refresh rate, and can be adjusted for participants with different inter-pupillary distances. Some examples of two-dimensional screen captures from the virtual environment are shown in **Figure 3**.

**Figure 3.**
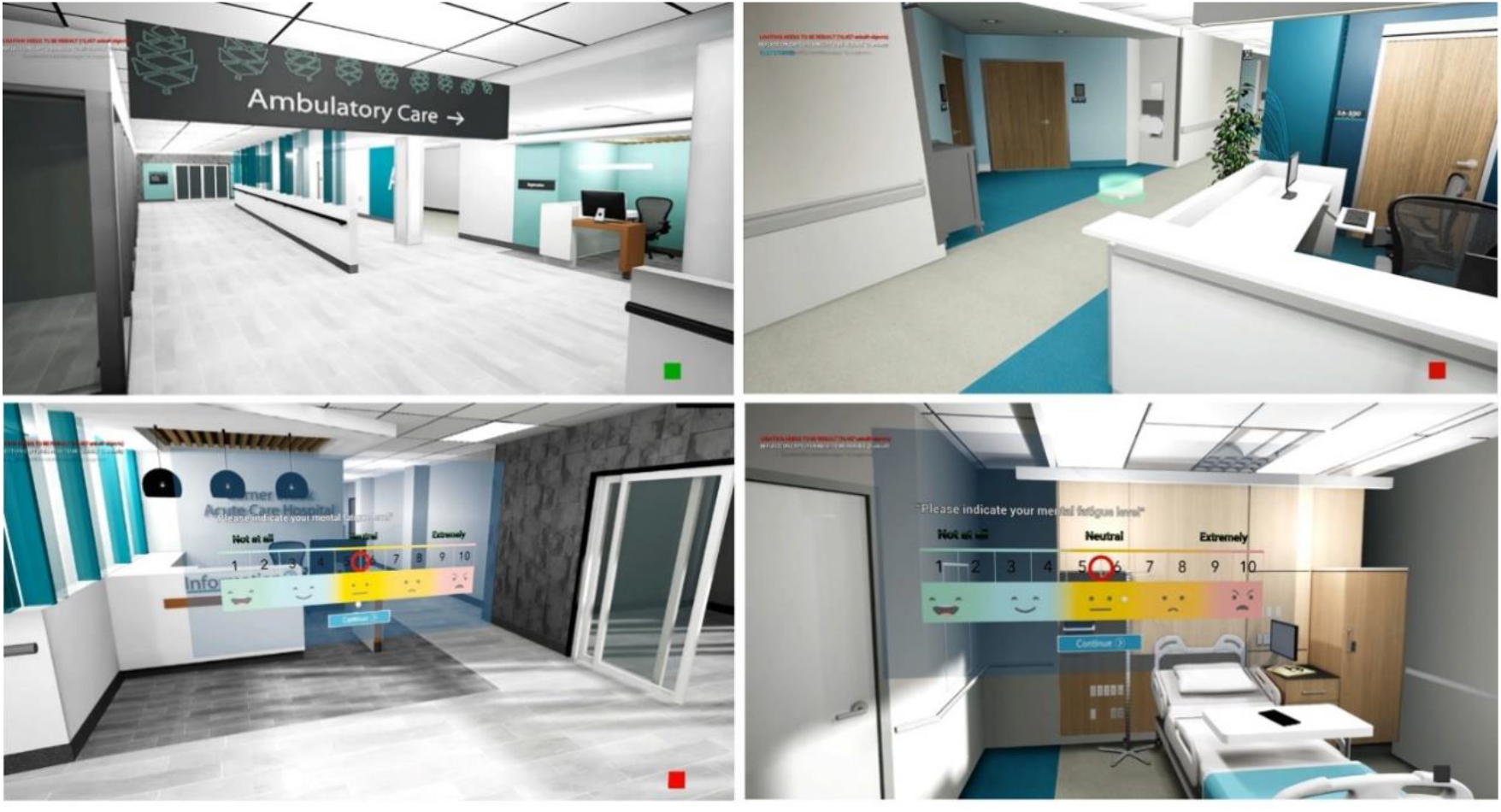
Screenshots from the participants’ view in the virtual environment. The bottom two images show our integrated Likert-scale widgets that simplify the collection of subjective feedback. The integrated software platform also helps to keep track of the duration of the wayfinding tasks, the position of the user in the environment, and the times at which the user looks at certain signs or other selected environmental features.

### Participants

A total of 81 research participants were recruited using a convenience sampling method (word-of-mouth and announcements on departmental e-mail lists). Ten of the participants had to be excluded due to technical problems during the wayfinding tasks, and an additional 8 participants had to be excluded due to technical problems in the data-recording (missing/incomplete data). The remaining 63 participants ranged in age from 18 to 55 years (*M*=20.72, *SD*=4.57). The majority were undergraduate university students (n=52), with a smaller number of graduate students (n=8) and faculty members (n=3). Demographically, 38 participants reported as female, 23 reported as male, and 2 preferred not to answer; and 26 participants reported as Asian, 8 as Latinx or Hispanic, 1 as Black, and 28 as White. All of the participants were associated with the Cornell University, representing the departments of Design, Psychology, Computer Science, Music, Biology, Marketing, Communications, Architecture, Human Development, Policy Analysis and Management, Development Sociology, Hotel Administration, Economics, Mechanical Engineering, Food Science, City Planning, Nutritional Sciences, and Chemistry.

We collected a variety of background data from the participants, which were not analyzed as prospective moderator variables but rather included here as an overall snapshot of the study sample. A majority of the participants (n=43, 68%) reported that they had experienced a hospital environment within the previous six months, either as a patient or as a visitor. On a Likert scale (1=not at all; 10=extremely), participants expressed moderate familiarity with hospital environments (M = 5.14, ±2.59 S.D.), and indicated on average that such environments were moderately stressful and confusing (M = 4.83 ±2.34 S.D.). Post-experiment surveys indicated that the participants on average were reasonably comfortable when wearing the VR headset (*M*=3.22, *SD*=1.70; 1=comfortable, 10=uncomfortable) and the physiological sensors (*M*=2.96, *SD*=1.83; 1=comfortable, 10=uncomfortable). Participants also reported that they felt the VR environment was reasonably realistic in both the inpatient areas (*M*=6.17; *SD*=2.12; 1=not similar to reality, 10=very similar to reality) and the outpatient areas (*M* = 5.91; *SD* = 1.99; 1=not similar to reality, 10=very similar to reality). In regard to sleep levels, 35% percent reported getting eight or more hours of sleep the previous night, 46% reported six to seven hours of sleep, and 19% reported four to six hours of sleep. None of the participants were aware of having current neurological conditions, and none reported the recent use of psychoactive substances (excluding caffeine).

Each participant gave informed written consent before participating in the experiment, and the overall study protocol was approved by the Institutional Review Board at Cornell University. All of the experiment sessions took place at the same physical location in Design and Augmented Intelligence Lab at Cornell. We used a randomized between-subject design, allocating each participant to one of the three condition groups in a pseudo-random order.

### Procedures

Sessions were conducted for one participant at a time. During each session, after providing consent the participant was carefully fitted with the physiological sensors by trained research team members. To establish resting-state data, after sensor fitting and quality checking, the participant was asked to sit quietly facing a blank computer monitor for one minute, and then to sit quietly with eyes closed for one minute. Once the resting-state data were collected, the participant was fitted with the VR headset and entered the virtual environment. An initial five-minute “free” period in the VR allowed the participant to become familiar with the navigational tools and to explore the platform. Two blocks of navigational tasks were then assigned, an “inpatient” block consisting of six wayfinding tasks within a virtual “inpatient” department, and a block of five wayfinding tasks in the virtual hospital’s “outpatient” area. All participants completed the same navigational tasks in the same sequence. To promote greater immersion, each task-series began with the presentation of a written scenario, asking the participant to imagine themselves in a moderately stressful medical situation (**Table 1 and Figure 4**).

**Table 1.** Wayfinding Task Descriptions

| **Inpatient Wayfinding Scenario** |  |
| --- | --- |
| Pre-task | “Imagine a beloved member of your family is going through a high-risk surgery, and you are late for the patient visit. You are in a big hospital and you want to visit your family member as quickly as possible. Please try your best to complete the following tasks as quickly as possible.” |
| Task 1 | Find the information desk. |
| Task 2 | Find the elevator. |
| Task 3 | Find the nurse station information desk in Ward A. |
| Task 4 | Find patient room #5A-511. |
| Task 5 | Find the elevator. |
| Task 6 | Find the main exit of the hospital. |
| **Outpatient Wayfinding Scenario** |  |
| Pre-task | “Imagine that you are feeling ill and need immediate treatment. You just want to find the fastest way to get to the room where you will complete your medical exams. Please try your best to complete the following tasks as quickly as possible.” |
| Task 6 | Find the ambulatory care reception desk. |
| Task 7 | Find treatment chair #4 in Section C. |
| Task 8 | Find the ambulatory care reception desk again. |
| Task 9 | Find the cafeteria cashier. |
| Task 10 | Find the main exit of the hospital. |

**Figure 4.**
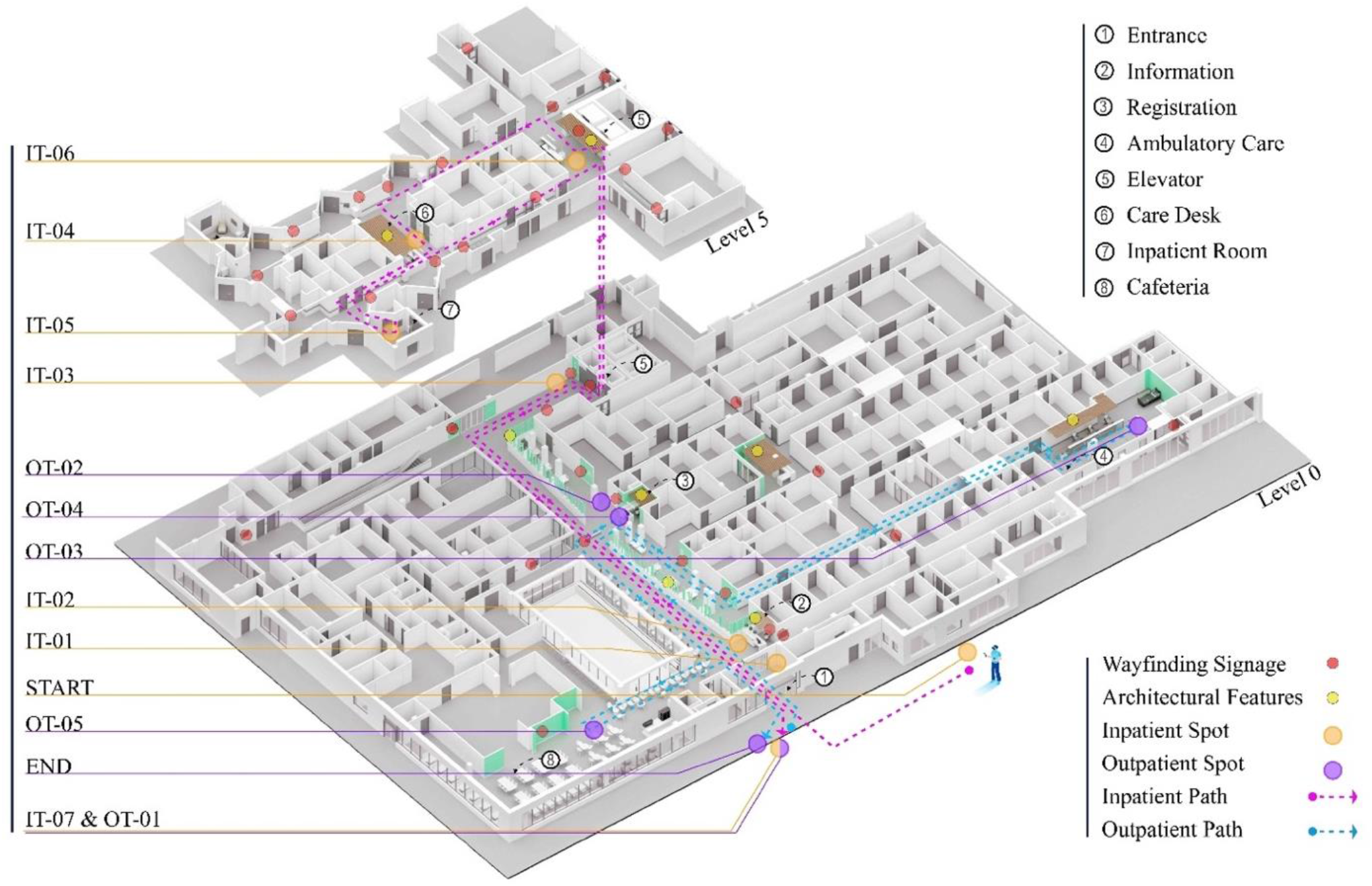
Navigational inpatient tasks (IT) and outpatient tasks (OT) as shown on hospital plans.

### Psychophysiological Data Collection

The sensors used in the study included a non-invasive EEG cap to record electrical brain activity, electrooculography sensors (EOG) to record eye motions, electrocardiogram sensors (ECG) to record heartbeat, a galvanic skin response (GSR) unit to record skin conductance, and a tri-axial head accelerometer to record head motions. All of the resulting physiological data were recorded at 512 Hz and synchronized using the 128-channel Actiview System (BioSemi Inc., Amsterdam, Netherlands) with Ag/AgCl active electrodes (**Figure 5**). The participant’s actions and motions within the virtual environment were integrated into the psychophysiological data using event markers manually inserted by trained research team members using millisecond accuracy achieved with Camtasia video-editing software (see ***Data Collection and Analysis***).

**Figure 5.**
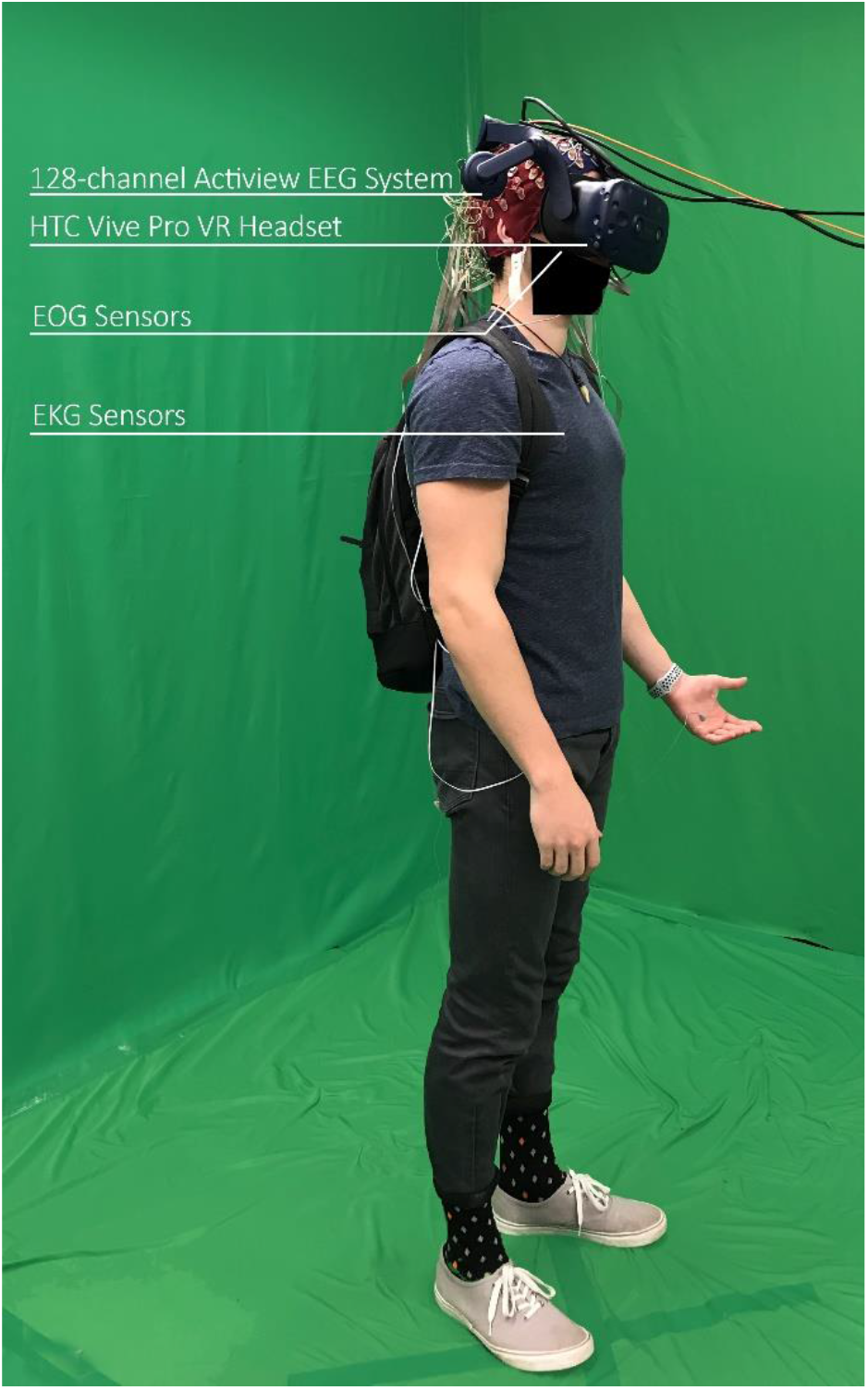
A study participant wearing the physiological sensors and VR display helmet.

Immediately after completing each navigational task, the participant responded to a quick set of Likert-scale questions (presented directly in the virtual environment) in order to self-report levels of mental fatigue, stress, and confusion during the preceding task. At the end of the entire session, the participant filled out an exit survey to provide additional feedback about the VR experience.

### Data Collection and Analysis

A custom event-synchronization technique was developed to help in synchronizing physiological, self-reported, and behavioral data as participants completed the wayfinding tasks. We included a small box in a corner of the VR environment, which is out of the participants’ field of view but can be seen in the observing monitor of the researcher’s computer. This box changes color according to events happening during the experiment, for example when the participant’s gaze crosses a designated signage area in close enough proximity for the sign to be legible. The screen on the researcher’s computer was recorded. This system helped the researchers to extract all the event markers from an Unreal Engine extension .log file to mark behavioral events as they occurred (sign views, direction changes, stationary periods, task completion, etc.). As a result, the event markers were created automatically based on a code developed by our research team to read the .log file, extracting time points for each event, and exporting into an MS Excel spreadsheet for further analysis. The time-stamps on these events can then be synchronized with the signal-derived physiological information, allowing the researchers to analyze physiological patterns that occurred during particular wayfinding experiences. After data collection, two researchers reviewed the recorded screen videos and added additional markers such as marking when participants are moving in the wrong direction or correct direction.

We established an exploratory analysis for session events, which included: (a) non-VR neural resting-state; (b) VR “free exploration” without tasks; (c) start of the navigation process; (d) successful ongoing navigation; (e) unsuccessful periods in the navigation process (e.g., being lost, confused, or stationary); (f) examining signs and other wayfinding markers; and (g) achieving the target destination. The amount of time that the participants spent in various states—particularly the unsuccessful navigation periods, time spent looking at signs, and overall task duration—were used as behavioral metrics in evaluating wayfinding success. Self-reported data in terms of fatigue level, stress level, and confusion level were analyzed separately. All of these responses were averaged across the participants in each design condition (A, B, and C), and then statistically compared to look for significant differences between the design conditions.

### EEG Preprocessing

The physiological data analysis in the current study focused on the EEG signals that were collected from the participants (the additional ECG, EOG, and GSR data will be triangulated in future work). The EEG data were analyzed using the EEGLAB software package (Delorme & Makeig, 2004). Raw “bdf” data files were imported at 512 Hz and downsampled to 256 Hz. The data were low-pass filtered at 100 Hz, high-pass filtered at 0.1 Hz, and then run through the Cleanline algorithm (Mullen et al., 2013), which selectively filters out the 60 Hz power-line noise using an adaptive frequency-domain (multi-taper) regression technique, to avoid distorting the data. The data was then run through the PREP Pipeline (Bigdely-Shamlo et al., 2015), which is a robust re-referencing method that minimizes the bias introduced by referencing using noisy channels. In addition to visual inspection for high-noise channels, bad channels were removed if they presented a flatline for at least 5 seconds, and if the correlation with other channels was less than 0.7. Time windows that exceeded 20 standard deviations were reconstructed using artifact subspace reconstruction (Mullen et al., 2013). Removed channels were reconstructed with spherical spline interpolation from neighboring channels. The data was re-referenced again to the average of all the 128 channels.

To compare between the three different design conditions (H3, H4), the pre-processed EEG data were epoched into 5s time-windows focused on the identified periods when each participant was viewing a navigational sign in the hospital VR environment, with each epoch starting 1 second before a sign was observed, and extending 4 seconds after the onset of sign-viewing. Each of these time windows was considered a trial. Given the nature of the experiment, with freely-behaving participants navigating through the environment to complete their tasks, the number of trials per participant varied (M=12.5 trials +/- 4.4 S.D.).

Statistically independent components of the EEG data were extracted using Independent Component Analysis (ICA), specifically the “extended Infomax” variation as implemented in the EEGLAB package. Independent Component Analysis (ICA) was conducted in order to isolate underlying neural sources, which are otherwise inseparably mixed via volume conduction (i.e. the spreading of electrical signals through any conductive space, aka the skull and scalp) at the EEG electrode locations (Onton et al., 2006). The independent components (ICs) were projected as equivalent dipoles that model the components’ source within the brain. Due to the prior removal of artifact-laden channels, the principal component analysis option was used in this analysis to reduce the data’s dimensionality and compute a number of ICs corresponding to the data rank (i.e. the number of non-interpolated channels). The equivalent dipole source-localization for the ICs was approximated using the EEGLAB dipfit package (Oostenveld & Oostendorp, 2002), with the source-space warped to fit a template MNI coordinate system (Montreal Neurological Institute, MNI, Montreal, QC, Canada). ICs with a residual variance of less than 40% were kept for further analysis. ICs with similar properties were clustered together across participants and compared between conditions, as described below.

### Clustering of EEG Independent Components (ICs)

The features of the independent component dipoles were compressed into 10 principal components (as specified here) and the ICs were separated into 16 clusters within EEGLAB using k-means clustering on weighted features: scalp topography (weight = 1), dipole locations (weight = 6), mean log frequency spectrum (weight=1), and event-related spectral perturbation (ERSP) (weight = 3). Cluster number (N) is chosen *a priori* and values of N between 6 and 20 tend to yield the best range of clusters, for datasets featuring 128 channels, such that at least several clusters recruit ICs from a majority of subjects. For clustering purposes, the frequency spectrum was obtained with the Multitaper power spectral density estimate (Thomson, 1982), with a focus on the range from 3 to 25 Hz. The ERSP was computed over 1 to 50 Hz in a logarithmic scale, using a wavelet transformation with 3 cycles for the lowest frequency and a linear increase with a frequency of 0.5 cycles (Makeig, 2016). ICs further than three standard deviations from any of the centroids obtained were considered outliers and removed from the analysis.

To ensure the clustering replicability, we repeated the k-means algorithm 10,000 times, and we selected the final solution based on a cluster ranking system (Gramann et al., 2018). Since the k-means algorithm is initiated randomly, each clustering iteration yields a solution. The total number of clusters was set to 16. First, the region of interest was defined as the Occipital cortex, Brodmann Area 18, with Talairach coordinates [x = –20, y = –90, z = 7]. Second, we calculated the following criteria for each cluster: (i) the minimum number of participants per condition with an IC in the cluster, (ii) the total number of participants, (iii) the ratio of ICs/participant, (iv) the spread (average square distance) of the cluster centroid, (v) the mean residual variance of the fitted dipoles, and (vi) the distance of the cluster centroid to the Talairach coordinates specified for the ROI. We added the constraints of at least 8 participants per condition and a maximum Euclidian distance from the region of interest of 10 mm. These quality measures were weighted (i = 4, ii = 4, iii = –3, iv = –1, v = –2, vi = –4) and summed to rank the solutions. The highest-ranked solutions was selected as the final clustering solution.

A cluster in Brodman Area 18, the visual association area of the occipital cortex, was chosen as the region of interest (ROI), based on findings of relevant signal clusters during navigation tasks in previous experiments (Djebbara et al., 2019). Neural dynamics from ICs clustered in the this ROI were compared among conditions using event-related spectral perturbation (ERSP) on the event trials. Oscillatory neural activity in the theta band in the occipital cortex has been associated with navigation in mobile experiments when participants integrated visual, vestibular, and proprioception cues to perform wayfinding tasks (Do et al., 2020). Occipital cortex alpha suppression has also been previously observed in spatial learning for maintaining orientation in both passive (Chiu et al., 2012; Gramann et al., 2010; Lin et al., 2015; Plank et al., 2010) and active navigation tasks (Ehinger et al., 2014). Alpha band activity is often inversely related to task-related neural activity in most regions; therefore, alpha suppression can be interpreted as greater activity. Occipital cortex modulation in the beta band is consistent with cognitive workload (Botvinick et al., 2001; Cavanagh & Frank, 2014), arousal (Faller et al., 2019), and engagement in architectural spaces (Banaei et al., 2017; Vartanian et al., 2013, 2015)Thus, our analysis assumes that heightened activity in this brain region is indicative of more effective wayfinding activity.

### EEG Event-Related Spectral Perturbation (ERSP)

We examined the event-related spectral perturbation (ERSP) patterns from the resulting cluster of interest, calculated and averaged across all of the ICs in the cluster, for each condition. The time period from - 1000 ms to -400 ms before the onset of the “sign-seen” annotation was used as the baseline for each trial. A spectrogram of all single trials was computed for all IC activation time courses using the *newtimef* function of EEGLAB, from 0.5 Hz to 50 Hz in logarithmic scale, using a wavelet transformation with 3 cycles for the lowest frequency and a linear increase with a frequency of 0.5 cycles. Permutation tests (1,000 permutations) were computed to evaluate statistically significant differences in ERSP between the three design conditions, at a significance level of *p* < 0.01.

## RESULTS

### Self-reported Stress, Mental Fatigue, and Confusion (H1a & H2a)

At the end of each navigational task, participants were asked to rate their stress level, mental fatigue, and confusion during the task using 10-point Likert scales. For each of these categories (stress, fatigue, and confusion), the data were averaged across all participants and all wayfinding tasks in each design condition. One-way ANOVAs were then conducted to compare the different design conditions. No significant differences were found. As a secondary exploratory analysis, we examined the data separately for each navigational task (**Supplementary Table 1**). This evaluation turned up significant differences in Task 7 (in which stress and confusion were significantly higher in the A and B design condition compared to the C design condition), and in Task 8 (in which stress and mental fatigue were significantly higher in the A and B design condition compared to the C design condition). However, for the rest of the tasks, no significant differences were found. Overall, therefore, H1a and H2a were not supported.

### Time Required for Task Completion (H1b & H2b)

The time required for task completion was averaged across all participants and all tasks in each design condition. One-way ANOVAs were used to compare the different design conditions. No statistically significant differences were found. As a secondary analysis, the data were evaluated separately for each wayfinding task (**Supplementary Table 2)**. Only Task 7 showed a significant difference between the design conditions, with participants in condition C completing this single task more quickly than those in condition A and B. Based on these findings, H1b and H2b were not supported.

### Orientation Behavior and Correct Choice of Direction (H1c &H2c)

Five variables representing participants’ behavior were extracted from the wayfinding process, including (a) the number of times a participant started traveling in the wrong direction, (b) the total time spent traveling in the wrong direction, (c) the number of times the participant viewed a sign, (d) the total time spent viewing signs, and (e) the number of times participants traveled in a correct direction after viewing a sign. We also calculated a relative variable of (f) “sign efficacy” as the number of times participants traveled in the correct direction after viewing a sign divided by the number of times the participant viewed a sign. One-way ANOVAs were used to compare the different design conditions for each of these variables.

No significant differences were found for the overall number of times participants started traveling in the wrong direction, or for the total time spent traveling in the wrong direction. However, the number of times that participants traveled in the *correct* direction *after viewing a sign* was significantly different across the three conditions (*F* (2, 48) = 13.15, *p* < .01, η_*p*_ = .35). Post-hoc test results using the Tukey HSD test showed that participants in Condition C (*M*_*C*_ = 16.43, *SD*_*C*_ = 3.91) were significantly more likely to orient themselves in a correct direction after viewing a sign, compared to participants in Condition B (*M*_*B*_ = 11.40, *SD*_*B*_ = 3.90, *p* < .01) and Condition A (*M*_*A*_ = 10.70, *SD*_*A*_ = 2.86, *p* < .01). There were, however, no significant differences in this variable when comparing between Condition B and Condition A. This finding seems to suggest that enhanced color condition were not enough to increase participants’ wayfinding success after viewing a sign; but that enhanced color, graphics, and architectural features did improve the outcomes of navigational choices after participants stopped to read an enhanced sign.

The number of times that participants viewed signs was significantly different across the three design conditions (*F* (2, 48) = 7.78, *p* < .01, η_*p*_ = .24). Post-hoc test results using Tukey HSD test showed that participants in Condition C (*M*_*C*_ = 18.25, *SD*_*C*_ = 3.87) viewed significantly more signs compared to those in Condition B (*M*_*B*_ = 13.86, *SD*_*B*_ = 5.05, *p* < .05) and Condition A (*M*_*A*_ = 13.15, *SD*_*A*_ = 3.31, *p* < .01). However, a comparison of the latter two conditions showed no significant difference. In a similar fashion, one-way ANOVA results showed the total time participants spent viewing signs was significantly different across three condition (*F* (2, 48) = 3.96, *p* < .05, η_*p*_ = .14). Post-hoc test results using Tukey HSD test showed significantly higher total time in Condition C (*M*_*C*_ = 40.33, *SD*_*C*_ = 13.86) compared to Condition A (*M*_*A*_ = 27.41, *SD*_*A*_ = 11.88, *p* < .05). However, comparisons of sign-viewing time between Condition C vs. Condition B and Condition B vs. Condition A showed no significant differences. Overall, these results indicate that enhanced color, graphics, and architectural features (Condition C) were important in drawing building users’ attention to navigational signs, and that better color and graphics alone did not accomplish this purpose.

Finally, the analysis of the “sign efficacy” variable using the one-way ANOVA test showed significant differences across the three conditions (*F* (2, 48) = 3.50, *p* < .05, η_*p*_ = .12). Post-hoc Tukey HSD test indicated that the wayfinding signs in Condition C (*M*_*C*_ = 89.96, *SD*_*C*_ = 9.21) were significantly more efficient compared to Condition A (*M*_*A*_ = 81.72, *SD*_*A*_ = 11.43, *p* < .05). Comparisons of Condition C vs. Condition B and Condition B vs. Condition A showed no significant differences. These results again indicate that the enhanced color, graphics, and architectural features added in Condition C helped to improve navigational effectiveness in a way that better color alone did not accomplish. Overall, these results indicate that H1c was not supported, but that H2c was supported (**Figure 6 and Supplementary Table 3**).

**Figure 6.**
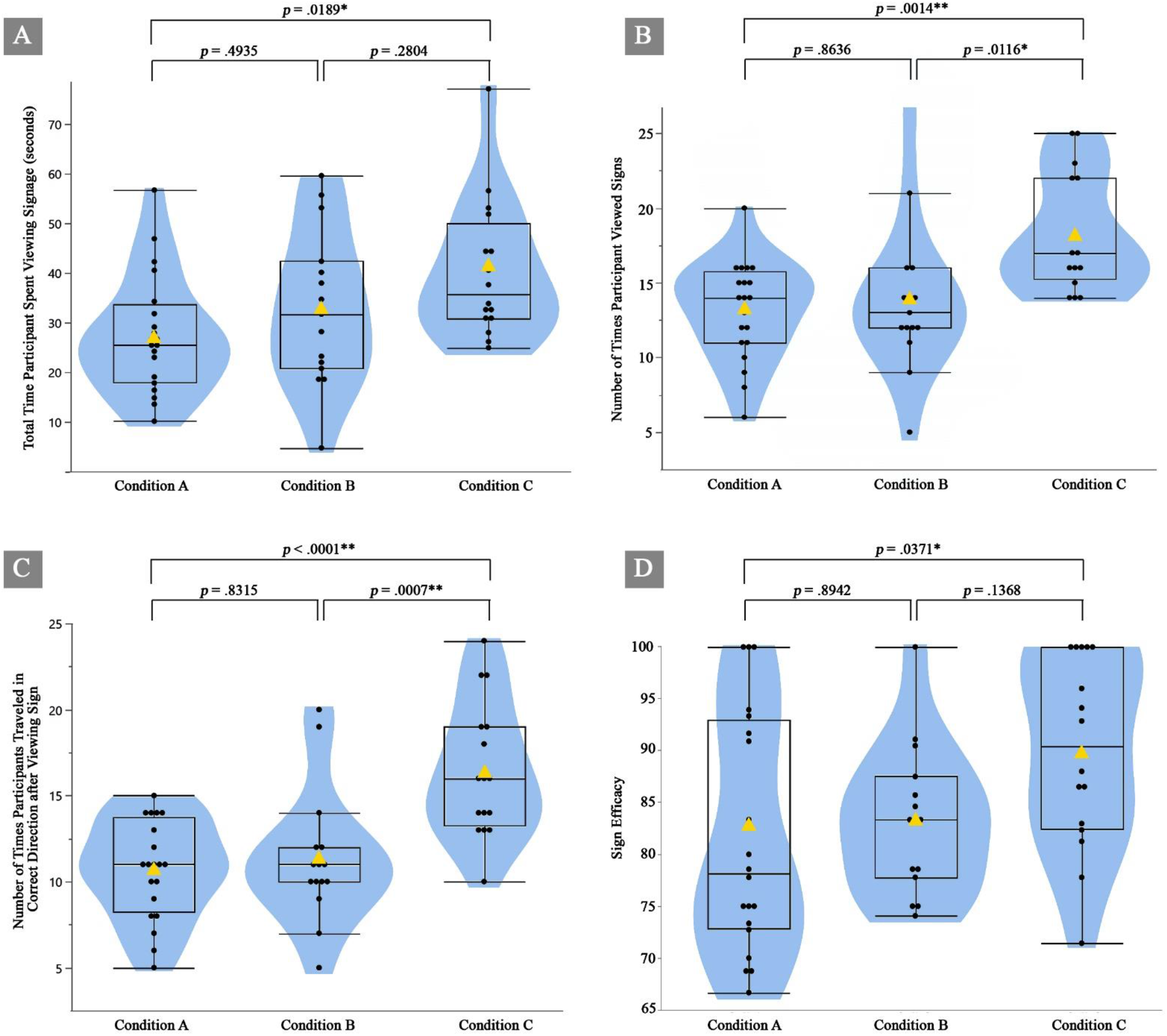
Results of Tukey HSD tests for (a) the amount of time participant spent viewing signs, (b) the number of times participants viewed signs, (c) the number of times participants traveled in the correct direction after viewing signs, and (d) the “sign efficacy” (number of times participants traveled in the correct direction after viewing a sign divided by the number of times the participant viewed a sign). Design Condition C showed significant advantages over the baseline Condition A in all of these categories, while Design Condition B did not. Yellow triangles show the mean. \**p* < .05. \*\**p* < .01.

### Neural Signatures of Spatial Awareness, Recall, and Cognitive Engagement (H3 & H4)

To test hypotheses 3 and 4, as mentioned above five-second EEG data epochs from 1 second prior to 4 seconds after observing wayfinding signs were extracted from all participants’ data. The neural dynamics of the most significant independent component (IC) cluster, which was localized to the visual association area Brodmann Area 18, were compared between the different design conditions across all participants.

#### EEG equivalent dipole cluster results

The optimal IC cluster that we identified was located with a centroid at Talairach coordinates [x = -21.7, y = -88.7, z = 10.6], in Brodmann Area 18. This cluster was seen in at least 8 participants per design condition, in 31 total participants, with 1.93 ICs per participant, 28.25 mm of average distance to the centroid, and 22% average residual variance. It was located 4.22 mm from the pre-defined region of interest (**Figure 7c**). The positive value-sided skewness of the weighed sum score (**7b**) from the individual criteria (**Figure 7a**) indicates that the automatic clustering algorithm found consistently at least one IC cluster around the original ROI. The optimal cluster selection only represents a marginal increase in the specified criteria, further validating the robustness of the IC characteristics found around the ROI.

**Figure 7.**
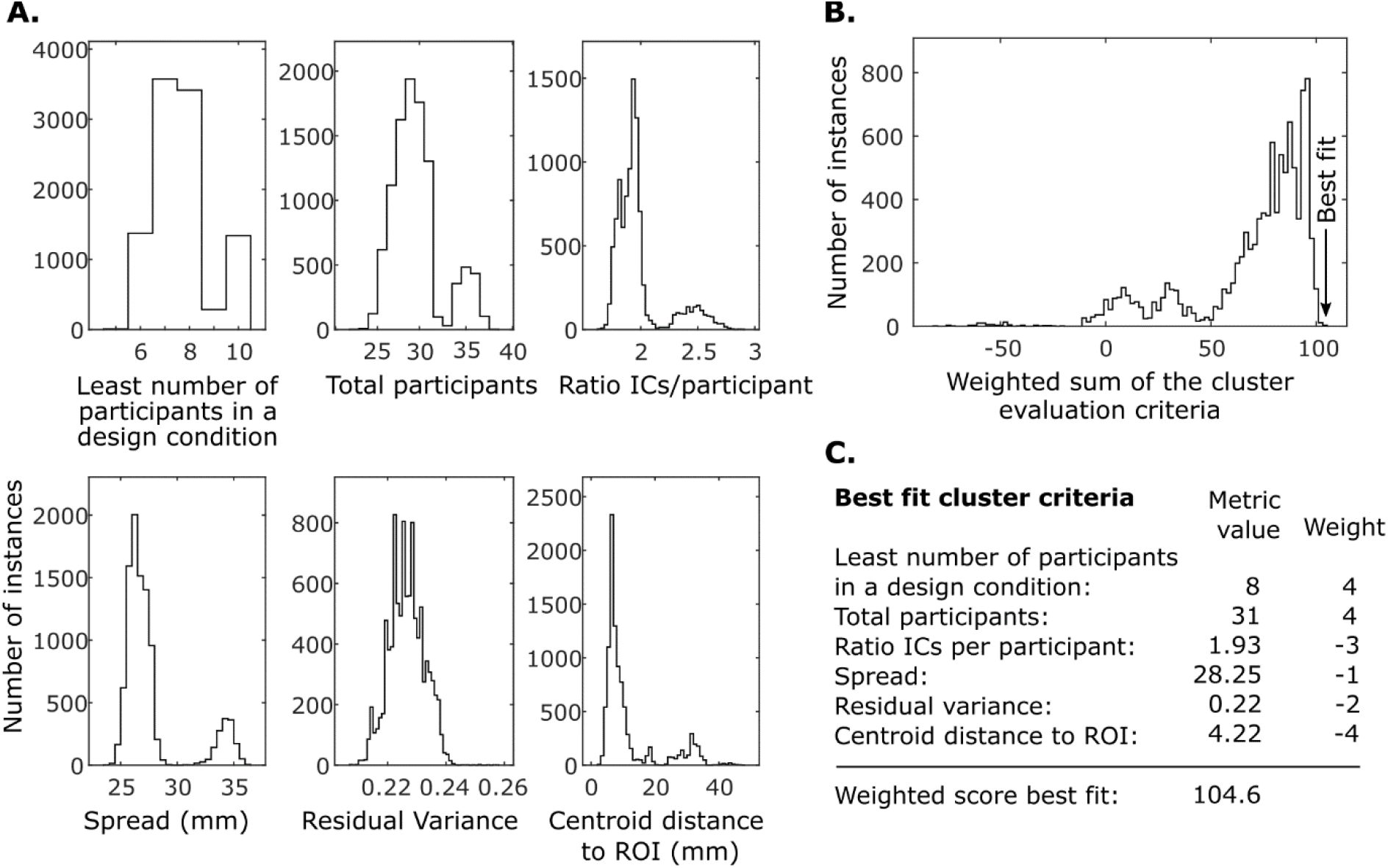
The EEG independent component cluster selection was based on six weighted criteria. The images here show (a) the distribution of cluster selection criteria among 10,000 random initializations, (b) distribution of the weighted sum of the six cluster evaluation criteria, and (c) the score of the best fit cluster on the basis of the specified criteria and corresponding weights.

An important criterion in the selection of the optimal IC cluster was that it should be reflected in a high number of participants in each design condition. We set the minimum limit for this number to 8 participants per condition. The selected optimal cluster met this requirement as it was reflected in 14 participants from Condition A, 9 participants from Condition B, and 8 participants from Condition C (**Figure 8b)**. The power spectrum density in the selected cluster showed consistent broadband desynchronization (less power) in design conditions C and B, compared to the baseline design Condition A (**Figure 8a**). There was a statistically significant difference across all frequency bands between Condition A and Condition C; however, a significant difference was found only in the delta and theta bands when comparing Condition B with Condition C (**Figure 8c**).

**Figure 8.**
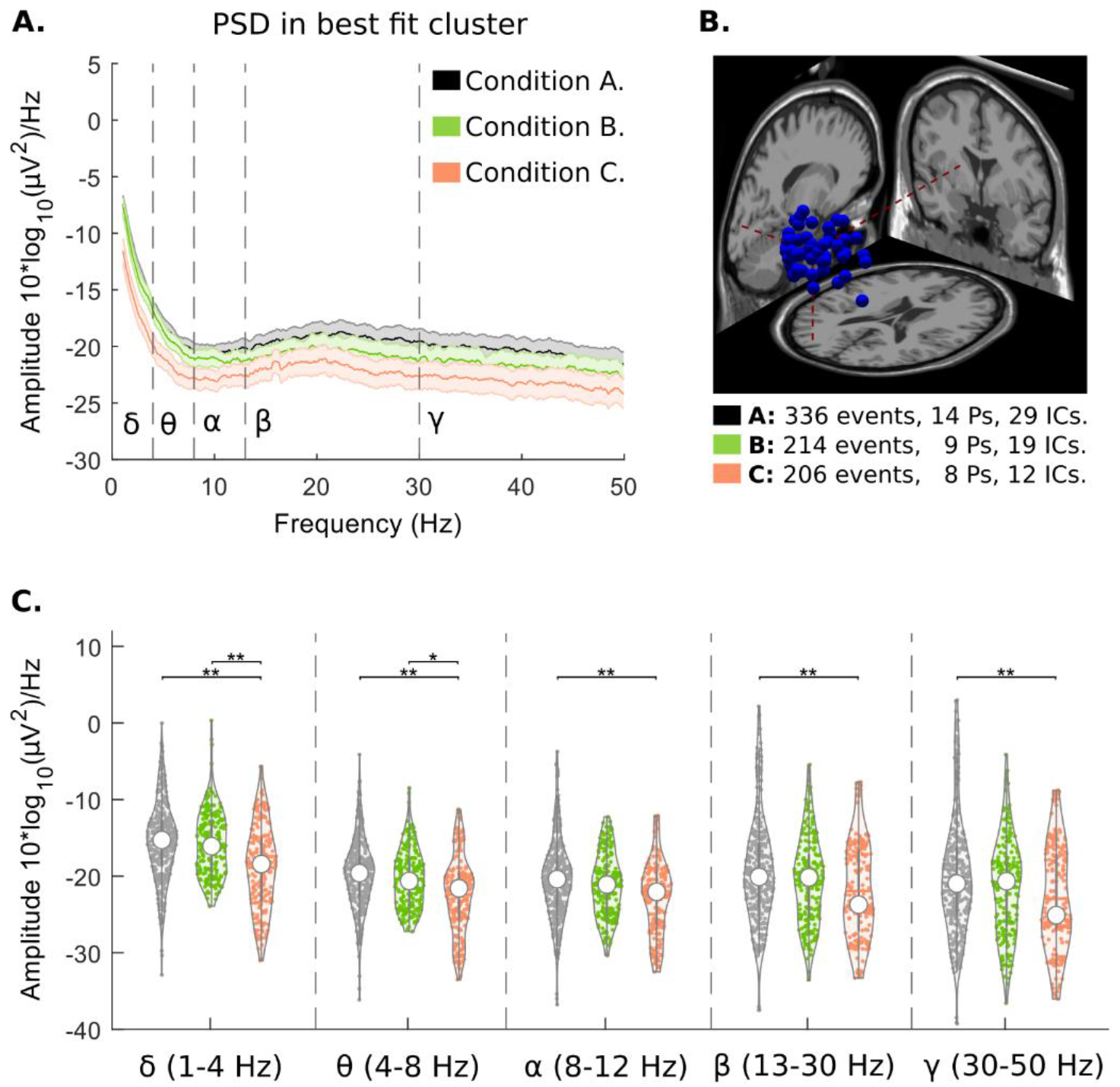
Attributes of the selected occipital lobe independent component cluster: (a) power spectral density with 99% confidence intervals, comparison between design conditions, (b) cluster dipoles and distribution among participants, and (c) frequency band-power distribution, comparison between design conditions (Kruskal-Wallis statistical significance: ** p < 0.01, * p < 0.05).

#### EEG event-related spectral perturbation results

The temporal dynamics of neural features can provide additional information about wayfinding success. To examine changes in the EEG signals over time, the power spectrum dynamics of the selected IC cluster were analyzed through 5-second time epochs associated with gazing at wayfinding signs in the VR hospital environment. As shown in **Figure 9**, statistically significant broadband desynchronization was seen in the comparison between Condition B vs. the baseline Condition A (particularly in the frequency range of 25–35 Hz), and also in the comparison between Condition C vs. the baseline Condition A (particularly at 1–12 Hz and 25–35 Hz). The comparison between Condition C vs. Condition B shows a much smaller area, and lesser intensity, of statistically significant desynchronization, located mostly in the 1–12 Hz range.

**Figure 9.**
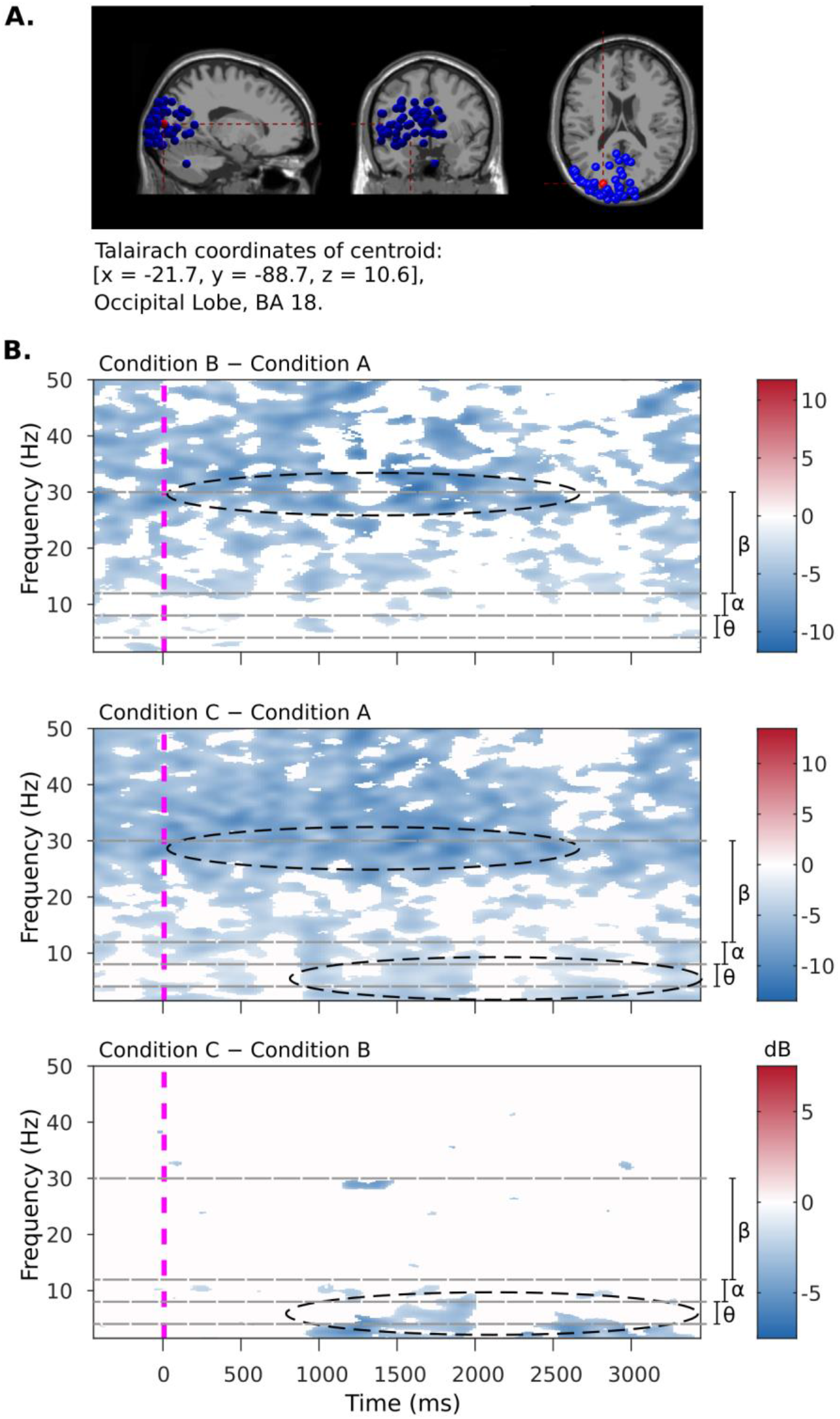
Event-related spectral perturbation from -1 to 4 seconds after a “wayfinding sign seen” event during the navigational tasks. The brain images (a) show the occipital cluster distribution of IC equivalent dipoles among participants (blue) and the cluster centroid (red) in the brain region Brodmann Area 18. The graphs (b) show ERSP comparisons between the three design conditions. The dotted ovals indicate comparisons with the largest effect size. White-colored areas show non-statistically significant effects, at a p-value < 0.01.

The analysis of design Condition C, when compared against both conditions A and B, shows a pattern of theta-band desynchronization that begins about 1s after looking at a wayfinding sign, continues until about the 2s mark, then disappears for about half a second, before returning again at 2.5s.

The EEG ERSP comparisons between conditions show statistically significant differences in the intensity of neural responses between the three design conditions, supporting both H3 and H4. These differences occur in the occipital cortex in the form of beta-band desynchronization between Condition A and B (H3), and additional oscillating theta-band desynchronization between Condition A and C (H4).

## DISCUSSION

The findings in this study regarding self-reported and behavioral data were mostly inconclusive, with a few notable exceptions. The participants did not report any significant differences in their stress levels, mental fatigue levels, or confusion levels after completing wayfinding tasks in the different hospital designs. There were also no significant differences in the time that was required for them to complete the wayfinding tasks. The data did indicate, however, that some orientation behaviors when viewing signs were more effective in design Condition C (enhanced color, graphics, and architectural feature) compared to Condition B (enhanced color) and Condition A (baseline). The participants who completed navigational tasks in Condition C looked at navigational signs more often and for longer periods of time, and were more likely to proceed in the correct direction after viewing a sign, compared to those who navigated through the other two hospital designs. While there was no significant difference in task completion and self-report stress, fatigue, and confusion in all wayfinding tasks, we found significant differences in these variables in some of the challenging outpatient tasks (Task 7 and 8). It is worth considering that nominal improvement would have long terms benefit with thousands of trips over a long-year life of a healthcare facility.

The EEG data taken from time-segments during which participants were viewing navigational signs further supported these behavioral findings. Frequency band-power analysis indicated significantly greater neural processing occurring when participants viewed signs in design Condition C and Condition B (**Figure 8)**, evidenced by beta (25-35 Hz) and theta-band (3-7 Hz) desynchronization compared to the baseline design condition. A temporal analysis of event-related spectral perturbations (ERSP) likewise showed significant desynchronization, indicating higher levels of cognitive engagement, when participants looked at signs in Conditions B to the baseline Condition A (**Figure 9)**, evidenced by sustained beta-band desynchronization. The ERSP analysis also showed a potentially oscillating behavior for the theta-band desynchronization in the seconds after participants viewed a navigational sign. Based on prior research (Rounds et al., 2020) this may indicate that the participants were cognitively integrating a broader array of environmental information after viewing the signs in design Condition C.

The theta-band desynchronization difference between Condition C and Condition B was found to be most prominent at 1s and 2.5 seconds after seeing the signage, sustained for approximately 1s. This observation suggests an oscillating form of theta-band desynchronization occurring when participants engage in wayfinding in a building with enhanced environmental affordances.

These EEG analyses focused on occipital and parietal brain regions that in prior research have been strongly associated with wayfinding and environmental-positioning cognition (Dhindsa et al., 2014; Lin et al., 2015; Rounds et al., 2020; Djebbara et al., 2020; Do et al., 2020). Based on these prior findings, we interpret beta and theta-band desynchronization in parietal and occipital brain regions as a likely indicator of integration of information from the environment for wayfinding; with a more pronounced effect when both architectural and signage enhancement (Condition C) compared to solely color enhancement in signage (Condition B) (**Figure 9**).

Our behavioral data in most cases only showed statistically significant differences between condition C and the rest of the conditions, while EEG analysis of beta and theta-band desynchronization in parietal and occipital brain regions yielded significant pairwise differences among all three conditions.

Taken together, these findings suggest that the enhancement of environmental affordances in the hospital design (adding architectural features and textures to highlight destinations and adding distinguishable patterns to signs) served to improve navigational cognition and orientation behaviors, in a way that enhanced color alone did not accomplish. In the enhanced color, graphics, and architectural feature condition the use of navigational signs was greater and more effective, and neural processes related to wayfinding were more active when viewing signs. These findings are generally commensurate with prior work in hospital wayfinding design (based mostly on anecdotal findings and static-image preferences) that have recommended incorporating both environmental affordances and manifest cues (Carpman, 1993; Devlin, 2014; Huelat, 2004; Marquardt, 2011; O’Neill, 1991; Passini et al., 2000; Pati et al., 2015; Rousek & Hallbeck, 2011; Ulrich et al., 2008).

The findings in our study indicated that overlaying indications of what should occur via color only provided little benefit, but this should not be interpreted to mean that they can be omitted. It is important to remember that design Condition C included *both* manifest cues and environmental affordances, and to recall that prior work (Vilar et al., 2014) has shown a heightened reliance on manifest cues in emergency situations.

It should also be reiterated that the improvements in orientation behaviors and cognition found in the study were not associated with significant improvements in the time that it took participants to complete the navigational tasks or improvements in their self-reported levels of stress, mental fatigue, and confusion. Further research is needed with additional facility designs and a broader participant base to determine whether or not this incongruity will be replicated. It could be that the difficulty of the wayfinding tasks in our experiment was not severe enough in relation to the capacities of our participant population to reveal significant effects in wayfinding outcomes, or that the design differences were not extensive enough to reveal such effects. Fortunately, due to the advantages of the VR testing platform, the possibilities are nearly endless for future testing of additional design variations and participant populations.

### Limitations

The use of virtual reality for design testing purposes allows for pre-construction testing of design effectiveness, and for rigorous comparative tests that would not be feasible to carry out in real-world environments. However, there are some limitations in the use of VR for behavioral studies, since the bodily motions, sensory immersion, and physiological responses to VR contexts may not precisely mirror real-world experiences. In the current study we encountered some episodes of “teleportation behavior” in the VR system (where participants would jump immediately from one position to another without a fluid transition). This glitch in the system’s realism may have impacted the wayfinding outcomes and the participants’ reaction to the environment. In a similar fashion, realism was reduced in the virtual environment due to the absence of any other humans in the hospital building. Typical hospital distractions that may affect wayfinding outcomes (noise and conversation, carts moving down the hallways, etc.) were not present. At the current time VR technologies are undergoing an extremely rapid pace of development, so researchers in this area should seek to stay abreast of new developments and take advantage of any improvements that can increase the level of realism and immersion for design-testing purposes.

The EEG analysis in this study was limited to a particular region of interest in the occipital cortex. Activity in this brain region has been robustly associated with wayfinding cognition, but it is not the *only* brain region involved in wayfinding. The process of parsing an environment, selecting navigational cues, making decisions, activating motion, and evaluating/correcting the results is neurologically complex. The selection of observed information during wayfinding can lead to different attentional mechanisms that elicit a motor response (Djebbara et al., 2020; Gallivan et al., 2018). The VR context, in which motion in the environment is initiated with hand-held controllers, may add further wrinkles to these neurological processes. In the current study we focused on visual attention (occipital cortex) as a correlate of motor execution action (Brown et al., 2011). This metric is well-supported for evaluating differences in wayfinding cognition, but a more robust neural picture could be obtained by investigating interplay between brain regions associated with sensory integration, planning, and decision-making (Cruz-Garza et al. 2020).

### Future directions

One near-future expansion of this research will involve the analysis of neural features during a wider array of wayfinding tasks. The current study only analyzed EEG data for brief time-periods after participants gazed at navigational signs. Additional design insight may be gained by looking at other wayfinding events, for example times when participants have taken a wrong direction and need to correct their wayfinding mistake. As noted above, the neurological analysis may also be expanded to additional brain regions, and it may be triangulated with additional physiological data such as heart rate or skin conductance (as measurements of stress).

One of the great advantages of this research platform is that it allows for the easy incorporation of new wayfinding design strategies and additional facility plans; expanding the number of designs tested is an important priority in generating more conclusive findings (Cruz-Garza et al. 2021). This is particularly exciting in that it provides an opportunity to test new and innovative wayfinding design strategies that might otherwise struggle to gain adoption. Expanding and broadening the study population is a priority as well—in the future we may be able to consider participant demographics (age, gender, health condition, familiarity with hospitals, etc.) as potential moderating variables.

### Implications for practice

The current study was undertaken in collaboration with Parkin Architects, and it evaluated real-world wayfinding designs in one of their ongoing healthcare facility projects. The outcome of this research includes specific findings about the proposed design of this facility; but perhaps more importantly, it also advances a new and tangible pre-construction testing method that can be directly applied to a wide variety of healthcare projects involving the evaluation or development of wayfinding systems as well as other aspects of healthcare planning and design. The testing and optimization of such designs prior to their implementation can prevent the need for costly changes after investment in physical construction.

On a broader level, the optimization of wayfinding systems can improve the experiences of patients, visitors, and staff members in healthcare settings, potentially reducing anxiety and ensuring that people get to their appointments on time (Davis & Ohman, 2016; Jamshidi & Pati, 2020; Mollerup, 2009; Rousek & Hallbeck, 2011). The provisional findings of this study suggest that environmental affordances, such as changes in material, lighting, and ceiling heights to highlight destinations and prompt accurate movements and adding distinguishable patterns to the wayfinding signs, may be a valuable aspect of wayfinding design. The study provides empirical evidence to support what has been an intuitive practice in wayfinding design in healthcare facilities. It will encourage the remaining skeptics or budget conscience owners/project managers to support or validate investment in enhanced environments for better wayfinding. The testing platform that we have created will lay the foundation for developing a larger body of knowledge in this area and conducting pre-construction design testing that would not otherwise be possible to carry out. Ultimately the use of rigorous virtual testing will help to promote the adoption of evidence-based principles in the industry, with the benefit of substantial cost savings and an improved environment for the public.

## CONCLUSION

This study applied a novel VR approach to evaluate wayfinding success under different architectural design conditions in a healthcare facility. Multiple data sources were collected and analyzed, including self-reported responses, behavioral metrics, and measurements of neural activity in wayfinding-relevant brain regions. Three variations of the interior hospital environment were created: a baseline design with standard signage and minimal environmental contrast (Condition A), a design featuring signage with added color and graphics to better highlight destinations and wayfinding information (Condition B), and a design that included the enhanced signage from Condition B along with improved architectural features and textures to assist in navigation (Condition C). Participants for the study were recruited using a convenience sample (mostly undergraduate students), were randomly assigned to one of the three design conditions, and were asked to complete various navigational tasks in the virtual hospital building.

The findings indicated that orientation behaviors and wayfinding cognition were significantly improved in Condition C, compared to the other two interior hospital designs. Few differences in wayfinding metrics were found between Condition B and Condition A, indicating that adding color to highlight destination and increase the contrast of wayfinding information against the overall environment alone may not be adequate to promote better wayfinding. These findings of the advantages in Condition C support the value of adding color combined with adding environmental affordance features in healthcare design for wayfinding purposes; examples of this approach include using different ceiling heights or materials to signal functional changes between different areas of the building, using lighting and textures to prompt or deter directional movement, and using distinguishable patterns to wayfinding signs. Since each facility design and user population is unique, caution should be used in generalizing the outcomes of this study to other healthcare settings. However, the virtual testing platform that we developed in this project can make it relatively easy for other researchers and designers to carry out their own studies of various under-development facility designs, which has the potential to lead to a larger corpus of findings and a deeper synthesis across them.

## ACKNOWLEDGEMENTS

The authors gratefully acknowledge Parkin Architects, the Government of Newfoundland and Labrador, the Western Regional Health Authority and the Corner Brook Acute Care Hospital. The authors thank the team at the Design and Augmented Intelligence Lab at Cornell University, including Julia Kan, Jeffrey Neo, Mi Rae Kim, Matthew Canabarro, Emme Wong, Clair Choi, Elita Gao, and Michael Darfler for assisting in data collection. The authors also thank the interior design and wayfinding/signage design team at Parkin Architects in Joint Venture with B+H Architects.

